# Vascular tropism models of blood-borne microbial dissemination

**DOI:** 10.1101/2021.05.05.442761

**Authors:** Anna E. Boczula, Amy Ly, Rhodaba Ebady, Janet Cho, Zoha Anjum, Nataliya Zlotnikov, Henrik Persson, Tanya Odisho, Craig A. Simmons, Tara J. Moriarty

## Abstract

Similar to circulating tumour and immune cells, many blood-borne microbes preferentially “home” to specific vascular sites and tissues during hematogenous dissemination ^1–5^. For many pathogens, the “postal codes” and mechanisms responsible for tissue-specific vascular tropism are unknown and have been challenging to unravel. Members of the Lyme disease *Borreliella burgdorferi* species complex infect a broad range of mammalian tissues and exhibit complex strain-, species- and host-specific tissue tropism patterns. Intravenous perfusion experiments and intravital microscopy studies suggest that heterogeneous tissue tropism properties may depend on tissue-specific differences in host and microbial molecules supporting vascular interaction and extravasation. However, interpreting these studies can be complicated because of the immune-protective moonlighting (multitasking) properties of many *B. burgdorferi* adhesins. Here, we investigated whether *B. burgdorferi* vascular interaction properties measured by live cell imaging and particle tracking in aorta, bladder, brain, joint and skin microvascular flow chamber models predict strain- and tissue-specific dissemination patterns *in vivo* These studies identified strain- and endothelial cell type-specific interaction properties that accurately predicted *in vivo* dissemination of *B. burgdorferi* to bladder, brain, joint and skin but not aorta, and indicated that dissemination mechanisms in all of these tissues are distinct. Thus, the ability to interact with vascular surfaces under physiological shear stress is a key determinant of tissue-specific tropism for Lyme disease bacteria. The methods and model systems reported here will be invaluable for identifying and characterizing the diverse, largely undefined molecules and mechanisms supporting dissemination of Lyme disease bacteria. These methods and models may be useful for studying tissue tropism and vascular dissemination mechanisms of other blood-borne microbes.

## MAIN TEXT

### Generation of hTERT-immortalized primary human endothelial cells (ECs) from tissues targeted by disseminating *B. burgdorferi*

One barrier for flow chamber studies of bacterial-endothelial cell (EC) interactions is the challenge of cultivating a sufficient number of primary human ECs, which undergo replicative senescence and phenotypic drift in culture. The first goal of our studies was to develop stable, replicable flow chamber models for studying bacterial-endothelial interaction mechanisms. Immortalization of ECs without loss of characteristic phenotypic properties has been achieved by retroviral expression of human Telomerase Reverse Transcriptase (hTERT) ^6^. We immortalized primary human ECs derived from umbilical vein (HUVEC), aorta, bladder, brain and joint synovial microvessels by retroviral hTERT expression (**Fig. 1**; **Fig. S1**). We attempted to immortalize cardiac, dermal, liver and lung microvascular ECs, but found that these cell types did not grow well, quickly lost their parent morphology in culture, or could not be immortalized (data not shown). hTERT expression successfully immortalized HUVEC, aorta, bladder, brain and joint ECs, as determined by comparing population doubling (PD) times of parental and hTERT-expressing progeny after 9-12 weeks of cultivation (**Fig. 1A**, **Fig S1A**).

**Fig 1:**
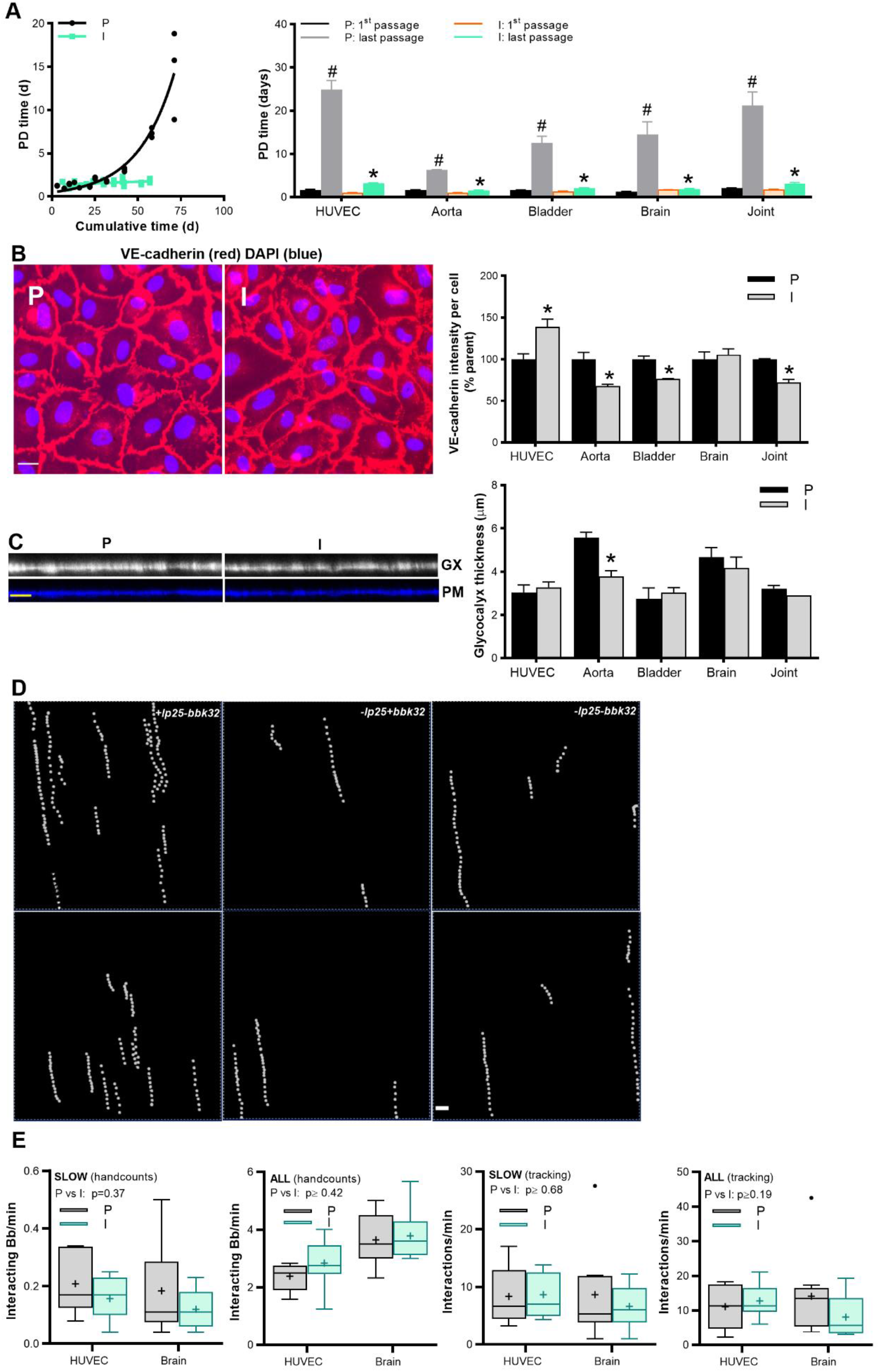
Generation and characterization of hTERT-immortalized primary human endothelial cells. Primary human endothelial cells (ECs) from vascular beds of indicated tissues immortalized by retrovirus-driven expression of human telomerase reverse transcriptase (hTERT). HUVEC are umbilical vein-derived ECs. **(A)** Sample non-linear regression showing change in population doubling time (PD) over time in culture for parent (P) and immortalized (I) brain microvascular ECs (left). PD plots and immunofluorescence (IF) images for other ECs are shown in **Fig. S1**. Mean ±SEM PD time at first and last passage for P and I ECs (right). **(B)** Representative epifluorescence IF photomicrographs of brain P and I ECs stained with antibody against endothelial marker VE-cadherin, and mean ±SEM VE-cadherin intensity/cell for all ECs. **(C)** Representative confocal 3D projection images of live brain ECs visualized under flow with fluorescent plasma membrane (PM) dye and wheat germ agglutinin (GX: glycocalyx), and mean ±SEM glycocalyx thickness for all ECs. **(D)** Sample 1 min time-lapse projection images of three genetically distinct bacterial strains (+lp25+*bbk32,* −lp25+*bbk32,* −lp25−*bbk32*) interacting with brain ECs under flow, captured by particle tracking. **(E)** Data are shown as Tukey box and whiskers plots of total (all) and slow (velocity <125 μm/s) interaction numbers under flow for all bacterial strains and HUVEC or brain ECs, measured by manual and particle tracking enumeration (hand counts, tracking, respectively). For all Fig. 1 experiments, N ≥ 3 independent EC and bacterial cultures. Statistics: two-way ANOVA of log-transformed values with Holm-Sidak post-tests. * indicates p < 0.05 P vs I (B, C, E, A: last passage). # indicates p < 0.05 1^st^ vs last passage for P or I ECs (A). Scale bars: 30 (B, C) and 20 (D) μm.

To determine if immortalized ECs retained primary EC properties we measured intercellular junction expression, localization of the EC marker VE-cadherin (**Fig. 1B**, **S1B**), and endothelial surface glycocalyx thickness (**Fig. 1C**) ^7,8^. Immortalization reduced VE-cadherin expression in aorta, bladder and joint ECs, but not HUVEC or brain ECs. (**Fig. 1B, S1B**). Glycocalyx thickness measured by live cell imaging under flow was similar in parent and immortalized progeny for all EC types except aorta (**Fig. 1C**), and was comparable to EC glycocalyx thickness measured by cryoelectron microscopy ^9^.

Finally, we determined if immortalization affected *B. burgdorferi*-endothelial interactions in flow chambers under postcapillary venule shear stress conditions at which *B. burgdorferi*-vascular interactions have previously been observed (1 dyn/cm^2^) ^10–15^. We used three genetically distinct infectious strains differing in carriage of plasmid lp25 or the gene encoding vascular adhesin BBK32 (+lp25+*bbk32*, −lp25+*bbk32*, −lp25−*bbk32*) ^10,11^. These strains exhibit distinct dermal postcapillary venule interaction properties in live mice which are recapitulated with HUVEC in flow chambers at 1 dyn/cm^2^ ^10–15^. We counted interactions manually and using particle-tracking methods (**Fig 1D**) to capture total (bacteria moving <175 μm/sec) and slow (bacteria moving <125 μm/sec) interactions for each strain and EC type. Manual counts measure the number of *B. burgdorferi* moving at these velocities over EC surfaces, whereas particle tracking captures numbers of binding-unbinding adhesion events that occur as bacteria move over surfaces at these velocities; each bacterial interaction trajectory consists of series of binding-unbinding events ^13^.

Immortalization did not affect *B. burgdorferi*-EC interactions, measured by manual or particle tracking, for EC types where VE-cadherin expression or glycocalyx thickness were reduced by immortalization (p≥0.42; **Fig. S1**: aorta, bladder, joint) and for EC types where immortalization did not disrupt these parameters (p≥0.19; **Fig. 1E**: brain, HUVEC). Although immortalization did not affect interaction numbers for aorta, bladder or joint ECs, we conducted subsequent analyses using data only from parent ECs for these cell types to ensure that unanticipated artefacts were not introduced, whereas analyses for HUVEC and brain ECs were conducted with both parent and immortalized cells.

### Genetically distinct *B. burgdorferi* strains exhibit different tissue tropism patterns following intravenous inoculation

Although it seems reasonable to assume that bacterial ability to interact with ECs under physiologically relevant fluid shear stress would affect their ability to disseminate out of the vasculature to specific tissues (tissue tropism), this hypothesis has not been formally tested. To test this hypothesis, we compared *in vitro* endothelial tropism patterns for different *B. burgdorferi* strains to *in vivo* tropism patterns in corresponding tissues measured in *in vivo* perfusion experiments (**Fig. 2, S2**). We used a short term intravenous (IV) perfusion model employed previously for tissue tropism and bloodstream survival studies ^16,17^, but adapted this model to permit inter-strain and inter-mouse comparisons independent of potential inoculation and immune clearance differences among C3H/HeN mice and bacterial strains. In the original model, blood is collected 1 h after inoculation of mice, followed by intracardiac saline perfusion to remove unattached bacteria, and quantitative real-time PCR (qPCR) to measure bacterial burden. To control for potential clearance and inoculation differences among bacterial strains and mice, we expressed dissemination to each tissue in individual mice as a percentage of total dissemination to all tissues measured in the same animal. Tissues examined were bladder, brain, heart, patella and skin, reflecting the EC types used in flow chamber experiments. Studies were performed at one hour, one day and one week after inoculation in both male and female mice to capture dissemination to all tissues independent of potential differences in dissemination kinetics, and to determine if sex affected outcomes. HUVEC has already been established as an effective model of *B. burgdorferi* murine dermal microvascular interactions ^13–15^. The aorta and brain human ECs used in our studies were previously found to have similar characteristics as corresponding EC types in mice ^18–22^. To the best of our knowledge, comparisons of primary bladder and joint human and mouse ECs have not been reported.

**Fig 2:**
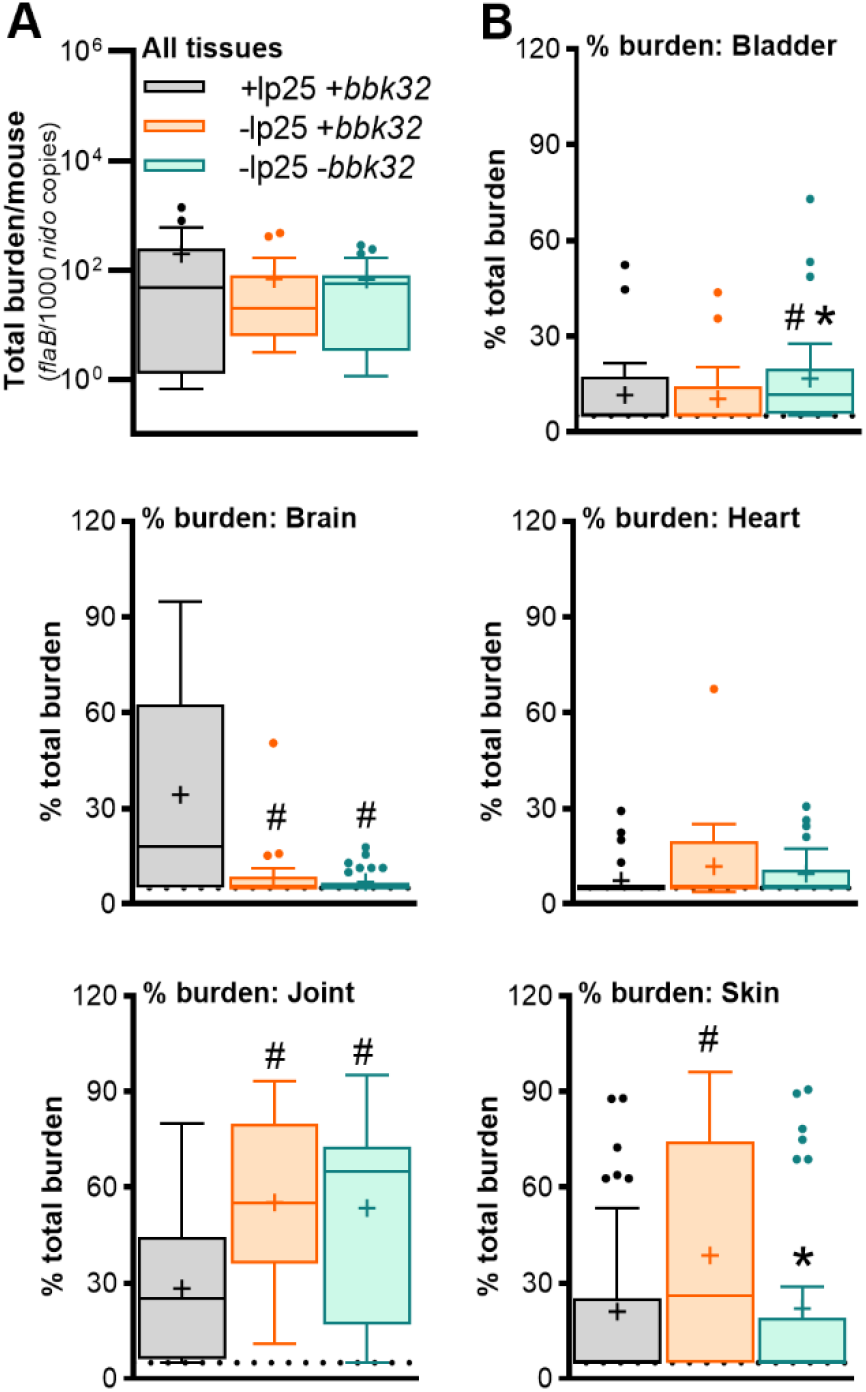
Genetically distinct *B. burgdorferi* strains exhibit different tissue tropism patterns following intravenous inoculation. Total *B. burgdorferi* burden in all tissues **(A)** and percent of total burden in indicated tissues after intravenous inoculation of male mice **(B)**, measured by quantitative PCR. Shown are values for all timepoints 1h, 1d, 1w. Values for individual timepoints for male and female mice are provided in Fig. S2. Data are depicted as Tukey box and whiskers plots. N≥28 mice/experimental group. Statistics: Kruskal-Wallis ANOVA of log-transformed values, with Dunn’s post-tests. * indicates p<0.05 vs −lp25−*bbk32*. # indicates p<0.05 vs +lp25+*bbk32*.

Unexpectedly, total burden across all tissues and at all timepoints was at least 1,000-fold greater in females than males (**Fig. S2A**). Although early strain specific dissemination patterns were similar in male and female mice, surprisingly we found that no strain disseminated appreciably or more than transiently to bladder or brain in female mice, whereas dissemination to heart was attenuated in male mice (**Fig. S2B**). Sex-specific *B. burgdorferi* tissue tropism patterns have not been systematically investigated, although we have previously noted sex-specific differences in brain and heart burden in long-term mouse infections with the −lp25*+bbk32* strain ^23^. Since we could not be certain that the dissemination patterns in females were not artefacts due to extremely high overall burdens, we conducted subsequent analysis using dissemination data from male mice (**Fig. 2**).

Consistent with earlier studies implicating BBK32 in dermal vascular interactions and dissemination to skin ^10,12,13,24^, BBK32-expressing strain −lp25*+bbk32* disseminated to skin more efficiently than BBK32-deficient strain −lp25*+bbk32* **(Fig. 2B)**. Also as observed ^25^, BBK32 was not required for joint dissemination (**Fig. 2B**). Surprisingly, BBK32 expression in an lp25-deficient background (−lp25*+bbk32*) was associated with impaired dissemination to bladder (**Fig. 2B**). As reported previously for dermal postcapillary venule interactions, lp25 carriage suppressed skin dissemination^10^, but also suppressed dissemination to joint and promoted dissemination to brain (**Fig. 2B**). No-strain specific dissemination differences were observed in heart (**Fig. 2B**).

### Bacterial-endothelial interaction numbers in flow chambers predict dissemination patterns *in vivo*

We next investigated which, if any, interaction properties measured for a specific bacterial strain and human EC type in flow chambers were associated with/predictive of dissemination to the corresponding murine tissue in perfusion studies (**Fig. 3, S1; Table 1**). *B. burgdorferi* interacts with ECs under flow conditions through transient interactions called dragging and tethering (**Fig. 3A**), as well as by stationary adhesion ^11^. Although stationary adhesion was once hypothesized to be the interaction step preceding *B. burgdorferi* extravasation, extravasation has only been directly observed following transient interactions ^11^. We found that stationary adhesion on ECs in flow chambers did not correlate with dissemination to any tissue examined (**Table 1**), implying that stationary adhesion is likely not a major determinant of tissue-specific dissemination.

**Table 1:** Correlation analysis, tissue burden *in vivo* (male, 1h, 1d, 1w post IV injection) vs flow chamber interaction numbers.

| **Parameter<sup>a</sup>** | **Aorta/Heart**<br>**(r, p)<sup>b</sup>** | **Bladder**<br>**(r, p)** | **Brain**<br>**(r, p)** | **Joint**<br>**(r, p)** | **HUVEC/Skin**<br>**(r, p)** |
| --- | --- | --- | --- | --- | --- |
| *Dragging bacteria/min* | n/a<sup>c</sup> | >0.95, <0.05 | >0.95, <0.05 | 0.50, 0.67 | >0.95, <0.05 |
| *Tethering bacteria/min* | n/a | >0.95, <0.05 | 0.50, 0.67 | >0.95, <0.05 | 0.50, 0.67 |
| *Total/min* | n/a | >0.95, <0.05 | 0.50, 0.67 | >0.95, <0.05 | 0.50, 0.67 |
| *Stationary adhesion/min* | n/a | <0.05, >0.95 | -0.50, 0.67 | <0.05, >0.95 | <0.05, >0.95 |
| *Vel<175 All* | n/a | >0.95, <0.05 | >0.95, <0.05 | 0.50, 0.67 | >0.95, <0.05 |
| *Vel<175 -T* | n/a | >0.95, <0.05 | >0.95, <0.05 | 0.50, 0.67 | >0.95, <0.05 |
| *Vel<175 +T* | n/a | 0.50, 0.67 | >0.95, 0.18 | >0.95, <0.05 | <0.05, >0.95 |
| *Vel<125 All* | n/a | >0.95, <0.05 | >0.95, <0.05 | 0.50, 0.67 | >0.95, <0.05 |
| *Vel 125-175* | n/a | 0.50, 0.67 | >0.95, <0.05 | >0.95, <0.05 | -0.50, 0.67 |

<sup>a</sup> interaction parameters calculated from particle tracking data, organized by interaction type: all, -T (untethered), +T
(tethered). <sup>b</sup> Pearson correlation coefficients, r (strain fold differences in tissue-specific burden *in vivo* vs strain fold differences in EC-specific flow chamber interaction numbers) and correlation p-values are indicated above each interaction group. <sup>c</sup> n/a: values for all strains in heart *in vivo* are identical (male)

**Fig. 3:**
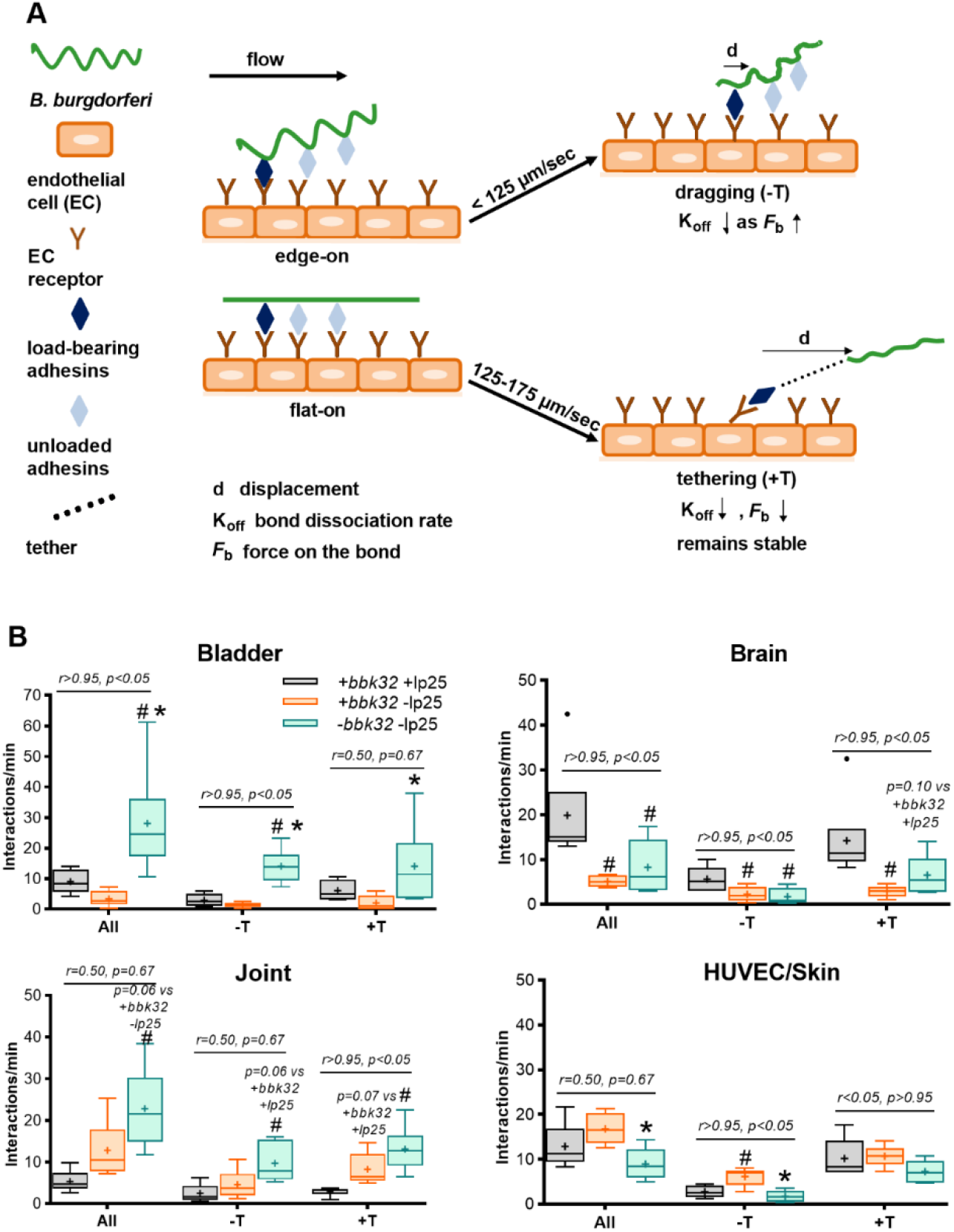
Bacterial-endothelial interaction numbers in flow chambers predict dissemination patterns *in vivo*. (**A**) Schematic depicting the two major modes of *B. burgdorferi*-EC interactions under physiological shear stress: Top: Slow, untethered (−T) “dragging” interactions in an edge-on orientation where mechanical load is transferred from peak to peak of the bacterial sine wave-shaped body (displacement, d, is wavelength), and that dissociate more slowly (K_off_) as force on the load-bearing adhesion bond (*F*_b_) increases. Bottom: Faster interactions in a flat orientation stabilized by tethers (+T) that slow dissociation by distributing mechanical load imposed on the adhesion bond; bacteria displace the length of the tether, which is further than peak-to-peak displacement in −T interactions. **(B)** *B. burgdorferi*-EC interaction number of interactions/min for all, −T (untethered) and +T (tethered) interactions in flow chambers, measured by particle tracking. Data shown as Tukey box and whiskers plots. N ≥3 biological replicates/experimental group. Statistical comparisons within each interaction group: ANOVA with Tukey’s post-tests. * indicates p<0.05 vs −lp25+*bbk32*. # indicates p<0.05 vs +lp25+*bbk32*. Pearson correlation coefficients, *r* (strain fold differences in tissue-specific burden *in vivo* vs strain fold differences in EC-specific flow chamber interaction numbers) and correlation *p*-values are indicated above each interaction group. Correlation values for all other interaction parameters are provided in **Tables 1, S1, Fig. S3**.

*B. burgdorferi* has a flat sine wave morphology and can interact with surfaces in flat or raised orientations (**Fig. 3A**) ^13,26^. Dragging *B. burgdorferi* moves slowly over EC surfaces (<125 μm/s) under flow, with the edge of its sine wave shape oriented perpendicular to the EC surface, and forms adhesion complexes that are not stabilized by tethers (**Fig. 3A** top: −T) ^13,14^. In this position, mechanical load is transferred from an adhesion complex at one “sine wave peak” to an adhesion complex on the next peak in the wave, and during each adhesion and load transfer event, the bacteria are displaced 2.83 μm, or the distance between sine wave peaks. These interactions depend on adhesion complexes that can sustain greater force on the load-bearing bond (*F*_b_) without dissociating and exhibit a smaller dissociation rate (K_off_). By contrast, tethering bacteria are oriented flat against EC surfaces under flow, displace further than 2.83 μm during each successive interaction, and move faster than 125 μm/s over EC surfaces (**Fig. 3A** bottom: +T) ^13,14^. Tethered interactions are stabilized by as yet unidentified tethers that absorb/reduce the force imposed on load-bearing adhesion complexes under flow.

To determine which, if any, interaction type or property predicted dissemination to corresponding tissues *in vivo*, we manually counted numbers of dragging and tethering bacteria and total interacting bacteria, and used particle tracking to identify and count total (Vel<175 All), tethering (Vel<175+T, Vel 125-175) and dragging interactions (Vel<125 All, Vel<175-T) (**Table 1**). We also investigated whether other biophysical interaction properties, including velocity, dissociation rate, displacement, force on the bond or number of successive interactions per trajectory were associated with tissue-specific *in vivo* dissemination patterns (**Table S1**; **Fig. S3**). To compare these variables, on widely different scales, to tissue burdens *in vivo*, we normalized the values for each parameter to the median value of the same parameter for the −lp25*-bbk32* strain, then performed correlation analysis. Since *B. burgdorferi*-HUVEC interaction properties in flow chambers correspond to interaction properties measured in the skin of live mice by IVM ^13–15^, we compared HUVEC interaction values to skin dissemination values *in vivo*. *In vivo* values used for comparison were pooled from all timepoints (**Fig. 2**). No significant correlations were observed for aorta ECs and heart, possibly because no significant strain differences were observed in hearts of male mice *in vivo* (**Table 1**) and possibly because mouse and human aorta ECs are subjected to much higher shear stress conditions *in vivo* (>10 dyn/cm^2^) ^27,28^ than the shear stress used in our studies (1 dyn/cm^2^).

Overall, the strongest predictors of strain- and tissue-specific dissemination patterns *in vivo* were numbers of interactions measured by manual or particle tracking enumeration (**Fig. 3B, Table 1**). Dissemination to bladder, brain and skin was associated most consistently with numbers of slow (velocity <125 μm/s), untethered (−T), dragging interactions observed on bladder and HUVEC ECs in flow chambers, whereas joint dissemination was associated most strongly with fast (velocity >125 μm/s), tethered (+T) interactions on joint ECs (**Table 1**). Dissemination patterns *in vivo* were not generally correlated with interaction velocity, dissociation rate, displacement, force on the bond or number of successive interactions per trajectory (**Table 1, S1, Fig. S3**), although it is possible that some of these properties might still be found to predict dissemination if flow chamber experiments are performed over a range of shear stress conditions that select for the most stable interactions ^13–15^.

These observations implied that in bladder, brain, joint and skin, dissemination out of the blood stream was limited mainly by strain- and EC type-specific availability of bacterial adhesins and cognate EC receptors supporting transient, dynamic adhesive interactions. That is, the rate-limiting step for interactions is bond association rate. We conclude that EC interaction numbers in flow chambers robustly predict dissemination to corresponding tissues *in vivo* when tissue burden is adjusted for mouse- and bacterial strain-specific differences in overall burden, and that *B. burgdorferi*-vascular interactions are likely a rate-controlling step of dissemination *in vivo*.

### Diverse tissue-specific vascular dissemination mechanisms

Consistent with previous sequencing studies of population bottlenecks during dissemination ^29^, we found that all strains could interact with all ECs and disseminate to all tissues to some degree (**Fig. 2, 3**), implying that multiple dissemination mechanisms exist for most vascular beds and tissues. *B. burgdorferi* produces at least 19 surface-exposed adhesins, 17 of which have been identified as possible or confirmed mediators of tissue-specific dissemination and/or colonization, and roughly half of which are known or predicted to moonlight in immune evasion ^3,17,30–39^. Strain- and species-specific polymorphisms in several *B. burgdorferi* adhesin genes also contribute to tissue-specific colonization patterns ^3,40^. Thus, dissection of tissue-specific dissemination mechanisms *in vivo* can be challenging. Quantitative, easily genetically and biochemically manipulated *in vitro* models that eliminate confounding factors such as immune clearance and extravascular conditions affecting microbe proliferation will be helpful for characterizing these mechanisms. However, despite the existence of multiple dissemination mechanisms for each tissue, we found that interaction/dissemination patterns for each bacterial strain, EC type and corresponding tissue were different, implying that distinct primary vascular interaction mechanisms likely support dissemination to each tissue (**Fig. 2, 3B**).

Bladder EC interactions and dissemination were lp25-independent and suppressed by BBK32 (**Fig. 2, 3B**). Candidate mediators might include glycosaminoglycan- and laminin-binding Lmp1 and BB0406, which have been linked to bladder in intravenous phage display and long-term infectivity studies ^41–45^. Since BBK32-expressing *B. burgdorferi* form a cap of polymerized plasma fibronectin at the mechanical load-bearing point of contact with ECs under flow ^14^, it is possible that BBK32 suppresses bladder dissemination by directly or indirectly blocking access of bladder-specific adhesins to cognate bladder EC receptors.

Joint EC interactions and dissemination were BBK32-independent, as previously reported ^16,25^, and suppressed by lp25 (**Fig. 2, 3B**). Candidate joint mediators identified previously by intravenous phage display, intravital microscopy and infectivity studies include Lmp1, BB0406, BB0347, OspC and VlsE, as well as P66, an integral membrane protein that mediates joint vascular transmigration but not EC surface interactions ^3,16,25,41,42,44,45^. These candidates bind a wide range of host molecules that could potentially be involved in joint vascular interactions and dissemination, including glycosaminoglycans, laminin, fibronectin, plasminogen and integrins. Of particular interest is OspC, which supports transmigration out of the joint microvasculature in a fashion that depends on its extracellular matrix-binding properties ^3^. It is not clear how or why joint EC interactions and dissemination would be suppressed by lp25, which also suppresses HUVEC interactions and skin dissemination (**Fig. 2, 3B**). We examined tissue-specific colonization data from the genome-wide signature-tagged mutagenesis library generated by Lin and colleagues ^45^, and found one lp25 locus (*bbe27*) where genetic disruption appeared to increase colonization of skin (ear) and joint compared to other tissues (bladder and heart) two weeks after infection. The *bbe27* open reading frames encodes conserved hypothetical protein of unknown function ^45^ that does not appear to be membrane localized (data not shown). It will be interesting to determine if this locus can affect EC interactions and intravenous tissue dissemination.

As reported previously ^10,12,13^, HUVEC interactions and skin dissemination were promoted by BBK32 and suppressed by lp25, but did not entirely depend on BBK32 (**Fig. 2, 3B**). Although *B. garinii* DbpA/B has been found to support HUVEC interactions in flow chambers, *B. burgdorferi* N40 DbpA/B does not ^46^, and the potential contributions of the *B. burgdorferi* B31 DbpA/B strain variant used in our studies to EC interactions and intravenous dissemination to skin have not been examined. Other candidates identified in intravenous infectivity studies are BB0406, P66 and the fibronectin-binding adhesin RevA ^16,44^.

Finally, we found that brain interactions and dissemination were BBK32-independent but promoted by lp25 (**Fig. 2, 3B**). Lp25 encodes the glycosaminoglycan-binding adhesin BptA, which is important for mouse infectivity in a tissue non-specific fashion (BptA-dependent brain colonization has not been examined) ^47^. Lp25 carries one additional open reading frame (*bbe09*) identified as a locus important for mouse infectivity in genome-wide transposon mutagenesis studies ^45^. The predicted gene product is a putative membrane-localized lipoprotein, P23, that is a member of the SP2/P23 conserved hypothetical protein family of unknown function ^45,48^. Another member of this family, the gene product of unknown function encoded by the *bbk52* locus on lp36, is transcriptionally upregulated in response to mammalian host conditions by the Rrp2, RpoN and RpoS regulators ^49^, and is transcriptionally upregulated in the central nervous system of non-human primates infected with *B. burgdorferi* ^50^. The potential contributions of BBE09, BBK52 and their encoding loci to *B. burgdorferi* pathogenesis have not yet been examined.

The ability to carefully dissect the molecular basis of tissue-specific vascular dissemination mechanisms using *in vivo*-validated model systems will be crucial for understanding the pathogenesis of *B. burgdorferi* and other blood-borne pathogens. These model systems also have strong potential for discovery of novel dissemination mechanisms using sequence-tagged random mutagenesis pathogen whole genome libraries and other, directed and undirected approaches.

## METHODS

### Ethics statement

This study was conducted in accordance with the most recent policies and Guide to the Care and Use of Experimental Animals by The Canadian Council on Animal Care. Animal work was approved by the University of Toronto Animal Care Committee in agreement with institutional guidelines (Protocols 20009347, 20009908 and 20010430). Work with *Borrelia burgdorferi* and primary human endothelial cells was approved by University of Toronto, Public Health Agency of Canada and Canadian Food Inspection Agency guidelines (University of Toronto biosafety permit 12a-M30-2).

### Mouse strains and husbandry

Intravenous perfusion experiments were performed using 6- to 7-week-old male C3H/HeN (Charles River Laboratories, Montréal, QC, Canada). Mice were housed in groups of 2 to 4 per environment-enriched cage with unlimited access to water and standard chow (Teklad 2018 Rodent Chow, Harlan Laboratories, Mississauga, ON, Canada).

### Primary human endothelial cell (EC) sources & cultivation

Specific endothelial cell (EC) types used in experiments are described in **Table S2**. All primary human ECs were purchased from Lonza (Lonza Inc. Allendale, NJ, USA) or Cell Systems (Cell Systems, Kirkland, WA, USA) and cultured at 37 °C according to the manufacturer’s instructions in tissue culture-treated T75 flasks without gelatin or Fn coating, in a humidified atmosphere containing 5% CO_2_. Lonza cells were cultured in EGM-2 medium (Lonza cat. CC-3156) supplemented with bullet kits CC-4147 (all cells except HUVEC) or CC-4175 (HUVEC). Cell Systems cells were initially thawed and grown for the first three passages in CSC medium supplemented with Culture Boost (Cell Systems cat. CSS-A104) and 5% heat-inactivated fetal bovine serum (FBS: Sigma-Aldrich Canada, Oakville, ON), then subsequently cultured in Lonza EGM-2 medium supplemented with bullet kit CC-4147. Cell Systems and Lonza cells were respectively frozen in Cell Freezing Medium (Cell Systems cat. 4Z0-705) or whole medium containing 1% dimethyl sulfoxide (DMSO; Bioshop Canada, Burlington, ON) and 20% heat-inactivated FBS. All ECs were passaged at ~80% confluence. Lonza cells, rinsed with 37 °C phosphate buffered saline without magnesium or calcium (PBS−/−: Multicell Wisent, St-Bruno, QC), trypsinized for 5 min at 37 °C with pre-warmed trypsin (0.05% trypsin/0.53 mM EDTA; Life Technologies Canada, Burlington, ON), followed by neutralization with complete medium. Cell Systems cells were rinsed with PRG solution (EDTA -dPBS, Cell Systems cat. 4Z0-610), trypsinized for 0.5-2 min (until rounded up but not detached) with 37 °C PRG-2 solution (Trypsin/EDTA -dPBS Solution cat. 4Z0-310), followed by neutralization with an equal volume of ice-cold PRG-3 solution (Trypsin Inhibitor-dPBS Solution cat. 4Z0-410). After trypsinization, all cells were centrifuged at 220 xg RT for 5 min followed by resuspension with fresh complete media and plated at 1:5 (Cell Systems cells)-1:10 (Lonza cells) dilutions. For all experiments only parent and immortalized ECs passaged fewer than eight times were used.

### Immortalization of primary human ECs

#### Retrovirus production

As previously described ^51^, primary ECs were immortalized by infection with retrovirus encoding hTERT, produced by transfection of Gryphon amphotropic packaging cells (Allele Biotechnology, San Diego, CA, USA) with plasmid pBABE-hygro-hTERT (cat. 1773; Addgene, Cambridge, MA, USA). Gryphon cells were cultivated according to manufacturer’s instructions at 5% CO_2_ 37 °C in 10 cm tissue culture-treated dishes (VWR Canada, Mississauga, ON) in DMEM-high glucose (Life Technologies) supplemented with 10% heat inactivated FBS and 100 U/ml penicillin/streptomycin (Thermo Fisher Scientific Canada, Burlington, ON). Gryphon cells were grown to 50-60% confluence by plating at 3.9-5.2 x 10^6^ cells/10 cm plate 18-24 h before transfection. On the day of transfection, 12 μl of FuGene HD transfection reagent (Promega, Madison, WI, USA) were added to RT serum-free Optimem medium (Fisher), followed by addition of 4 μg pBABE-hygro-hTERT prepared by maxiprep (Qiagen Canada, Mississauga, ON), to a final volume of 200 μl. This mixture was incubated for 15 min at RT, then added to Gryphon cells, followed by incubation at 32 °C/5% C0_2_ for ~24 h, when transfection medium was replaced with 10 ml fresh complete growth medium. Retrovirus-containing supernatant was harvested at 48 and 72 h after medium change, centrifuged at 2000 xg 4 °C 5 min to pellet cell debris, filter-sterilized with a 0.22 μm filter and stored in 2 ml single-use aliquots at −80 °C.

#### Retroviral infection and polyclonal selection

On the day of retroviral infection of ECs, 4.6 ml of 48 h viral supernatant mixed with 14.1 μl of 8 μg/ml polybrene (Sigma) was added to 1.4-2.8 x10^6^ ECs cultured to 20-30 % confluence in a T75 flask, followed by 3 h incubation at 37 °C then replacement of virus-containing medium with complete growth medium. This infection was repeated 24 h later using the 72 h viral supernatant. To select for polyclonal populations in which hTERT-expressing retrovirus was stably integrated, 48 h after the last infection hygromycin B (BioBasic Canada, Markham, ON) was added to endothelial cells to 20 μg/ml and included in culture medium for all subsequent passages ^52^. Hygromycin-resistant cells were frozen in whole medium containing 1% DMSO and 20% FBS upon reaching 80% confluence and at multiple passages thereafter.

#### Measurement of population doubling (PD) times

To determine if hygromycin-resistant cells were immortal, we measured PD times of triplicate cultures of parent and hTERT-expressing cells at each passage until ~2 weeks after parent cells reached replicative senescence (~2-3 months total passaging time) ^52^. Cells were counted at every passage (at 75-80% confluence) using a Beckman coulter Z1 particle counter (Beckman Coulter Canada, Mississauga, ON), blood cell counter vials (VWR) and 500 μl of a 10 ml resuspension of cells from each flask diluted with 19.5 ml Isoton II Diluent (Beckman). PD time, in hours, was calculated from the formula PD Time (PDT) = T*ln2/ln(Xe/Xb), where T is incubation time in hours. Xb and Xe are cell number at the beginning and end of incubation time, respectively. PD times were converted to days and plotted against cumulative days of cultivation using non-weighted least squares regression and exponential growth equations fit in GraphPad Prism v. 8.4.3 (GraphPad Software, San Diego, CA, USA).

#### Visualization and quantification of VE-cadherin-labelled EC junctions

ECs grown in triplicate to two days post-confluence in 48-well/9.8 mm plates (Falcon, Fisher) were rinsed with 37 °C PBS containing calcium and magnesium (PBS+/+: Sigma), fixed in −20 °C methanol at −20 °C for 15 min, rinsed with PBS −/− (Fisher), blocked for 20 min at 37 °C in PBS−/− with 3% w/v bovine serum albumin (BSA) (Sigma), incubated at 37 °C for 1 h with a 1:333 dilution of 1 mg/ml anti-VE-cadherin polyclonal antibody (Abcam, Toronto, ON, Canada, cat. ab33168) in PBS−/− with 3% BSA, washed once with PBS−/−, blocked for 30 min at RT with 10% heat-inactivated goat serum (Sigma) in PBS−/−, incubated in the dark for 1 h at RT with 5 μg/ml Alexa Fluor 488-conjugated goat anti-rabbit IgG (H+L) secondary antibody (Fisher) in PBS−/− with 10% goat serum, then washed three times with PBS−/−, followed by staining of cell nuclei for 3 min at RT with 1 μg/ml Hoechst 33342 (Sigma) in PBS−/−, and three final rinses with PBS−/−. Samples were stored in the dark at 4 °C until imaging.

Images were captured using an Olympus IX71 (Olympus, Tokyo, Japan) epifluorescence microscope equipped with a Retiga 2000R Fast-1394 camera (QImaging, Surrey, BC, Canada), X-Cite 120 series illumination source (Excelitas, Waltham, MA, US), DAPI and TXRed filters (Semrock Inc., Rochester, NY,US), a 40X (NA 0.4) air objective (Olympus, Tokyo, Japan) and and a custom acquisition script based on ImageJ (LOCI, University of Wisconsin, USA) ^53^ and Micro-Manager (Vale Lab, University of California, San Francisco, CA, US) ^54^ software. Image acquisition conditions were identical for all samples. Exposure times were 100 and 2000 ms for DAPI and TXRed, respectively, with no frame averaging, 0.75X zoom, 32 bits/pixel (1200×1600 pixels/image, 5.53 pixels/μm) and 0.18 μm^2^ xy pixel resolution.

Exported DAPI and TXRed 32-bit tiffs were converted to 8-bit in ImageJ, for all ECs, TXRed signal range was set to 200-800, and DAPI signal range was set to 200-800 (brain, HUVEC), 200-600 (aorta), 161-430 (bladder), 250-384 (joint).VE- cadherin signal intensity was measured in the TXReD images by subtraction of average background intensity (darkest 10×10 μm^2^ region of interest, ROI) from the average intensity measured in each of three 200×200 μm^2^ ROIs at three distinct locations in each image. The average of background-corrected signal intensity was measured per cell for the three ROIs in each image, and was expressed as a percent of the average intensity for parental EC images.

#### Visualization and quantification of EC glycocalyx thickness

As previously described ^13^, ~1.6 x 10^5^ ECs/channel were seeded in ibiTreat hydrophilic tissue-culture treated Ibidi μ-Slides VI^0.4^ (Ibidi, Planegg, Martinsried, Germany) and cultivated to 2 d post-confluence with daily feeding (a total of ~3 d). ECs were labelled for 10 min at 37 °C with 100 μl of a 1:10,000 dilution of Alexa 647-wheat germ agglutinin (lectin) (Life Technologies) to label the glycocalyx (GX) and for 3 min with CellMask Green plasma membrane live cell imaging dye (Life Technologies) prepared in endothelial growth medium to label ECs, then rinsed with perfusion medium before imaging. Perfusion media containing Hank’s Balanced Salt Solution (HBSS: Life Technologies), and 10 % heat-inactivated FBS (Sigma) was loaded in 60 ml syringes and perfused over endothelial monolayers in Ibidi chambers at 1.0 dyn/cm^2^ (34.1 ml/h) using a syringe pump (Model: NE1000, New Era Pump Systems Inc., Farmingdale, NY, USA). Ambient temperature on the microscope stage was maintained at 28 °C using an infrared heat lamp.

Z-series micrographs (512×512 pixels, zoom 1.25X, xy pixel size 0.73 μm, frame average 10, line average 3) were collected simultaneously every 0.75 μm in bidirectional resonant mode (pinhole 2.5 AU), at 511-600 nm (488 nm Argon laser, 10% intensity, HyD detector BrightR mode, gain 100), and 664-721 nm (633 nm laser, 100% intensity, HyD detector standard mode, gain 100), using an upright SP8 tandem scanner spectral confocal microscope equipped with a 25X 0.95 NA long working range water-immersion objective and operated by LAS X v.3.1.5.16308 software (Leica, Mannheim, Germany).

Glycocalyx thickness was measured using the line tool in Volocity v.6.5 (Quorum Technologies Inc., Puslinch, ON, Canada) at each of nine positions for each micrograph (xy pixel coordinates: 128×128; 128×256; 128×384; 256×128; 256×256; 256×384; 384×128; 384×256; 384×384), with a minimum of five independent biological replicates/experimental group. As described, ^55^ we first identified the positions of maximum intensity in each channel on the apical surface of endothelial cells. We calculated values for 25% maximum intensity, then, moving apically, identified the z-position at which the intensity in each channel reached 25% of the channel maximum. Glycocalyx thickness was calculated by subtracting the 25% intensity z-position in the plasma membrane channel (“outer edge” of plasma membrane) from the 25% intensity z-position in the glycocalyx channel (“outer edge” of glycocalyx). The median value from all nine measurements from the micrograph for each biological replicate (n≥5) was used for subsequent calculations and statistical analysis.

#### Preparation of endothelial monolayers for flow chamber experiments

ECs cultivated and seeded on Ibidi devices as described above were labelled for 5 min at 37 °C with 100 μl of a 1:2,000 dilution of CellMask Orange plasma membrane live cell imaging dye (Life Technologies) in endothelial growth medium, then rinsed with perfusion medium before imaging.

#### *B. burgdorferi* strains and preparation of bacteria for flow chamber experiments

All details of *B. burgdorferi* strains used in this study are described previously ^10–12^ and in **Table S4**. As reported ^13^ *B. burgdorferi* was grown, washed, resuspended to 4 x 10^8^/ml in ice cold PBS−/−, declumped, then diluted to 1 x 10^8^/ml in RT HBSS containing 10% heat-inactivated FBS immediately before imaging, except that bacteria were cultivated without addition of blood.

#### Live time lapse imaging conditions

Bacteria were perfused over endothelial monolayers in flow chambers as described in glycocalyx imaging section. For each biological replicate (independent endothelial and bacterial cultures) four 1-min xyt series (512×512, zoom 1.50X, xy pixel size 0.607 μm, line average 2) were acquired simultaneously in bidirectional resonant mode (pinhole 10.74 AU, frames per sec 14.08) at 497-535 nm (488 nm Argon laser, 90-98% intensity, HyD detector standard mode, gain 10) and 573-790 nm (561 nm laser, 9.8% intensity, PMT detector, gain 450-500) using the Leica upright SP8 tandem scanner spectral confocal microscope equipped with a 25X 0.95 NA long working range water-immersion objective. Time lapse videos were independently inspected by at least two individuals to confirm monolayer confluence and that at least 75% of the endothelial surface was at the imaging focal plane for at least 75% of video duration. Videos that did not meet these criteria were excluded from subsequent analysis. Subsequent quality control and analysis steps in the time lapse bioinformatics pipeline are summarized below and in **Fig. S4**.

#### Manual counting of *B. burgdorferi*-endothelial interactions under flow

Tethering (bacteria that pause but move faster than 125 μm/s) and dragging (bacteria that move <125 μm/s) interactions and stationary adhesions (bacteria that stay at the same position for ≥20 s) were manually enumerated in time lapse videos as described ^13^, except that dragging was counted in three 30×100 μm ROIs positioned at the centre of the field of view, and the average of this value was used for subsequent analyses. Analysis to identify high certainty outlier interaction values in each experimental group (typically 12-24 videos) was performed in GraphPad Prism (ROUT Q 0.1%). If outliers were identified in any video, it was removed from all subsequent analysis (**Fig. S4**). Interaction values for each biological replicate (independent bacterial and endothelial cultures) were calculated as the average of interaction values for all technical replicates (typically four videos). All subsequent statistical analysis was performed with values for biological replicates.

#### Particle tracking and measuring physical properties of *B. burgdorferi*-endothelial interactions under flow

Leica .lif library files were imported into Volocity, acquisition frame rates obtained from .lif files were entered manually, and the particle tracking was performed as reported ^13^, with the following modifications (bioinformatics pipeline summarized in **Fig. S4**). Each video was analyzed using six tracking protocols (**Table S4**), and interactions were identified in the resulting trajectories using a previously described formula spreadsheet that identifies individual interactions in trajectories (decelerating bacteria) and calculates physical properties of these interactions (see **Table S1** for parameters). To reduce potential user error in handling hundreds of thousands of lines of data from nearly 700 videos of data, macros were developed to permit batch data import into this formula spreadsheet without copying and pasting (**Fig. S4**). All parameter calculations and identification of untethered (−T) and tethered (+T) interactions were performed as reported ^13^, except that for *K*_*off*_ calculations time durations for interaction populations were binned in intervals of 0.07 s, and linear regressions and runs tests were performed in GraphPad Prism using only interactions with durations ≤0.24 s (>95% of interactions). R^2^ values for all curves were ≥ 0.98 and deviation from linearity was not significant (p>0.05).

Deduplication (arising from identification of the same trajectory with different tracking protocols) was performed by identifying interactions captured at the same time point in each video and examining their centroid xy position in the field of view. Interactions in the same quadrant were flagged as potential duplicates and manually verified in the original videos. Duplicate flagging using centroid position alone successfully identified duplicates in >95% of cases (verified by manual inspection). When duplicates were identified, the interaction where measured bacterial length was longest was retained for subsequent analyses, and all other interactions were removed (**Fig. S4**). After deduplication, interactions were then filtered to eliminate non-specific interactions with displacement ≥45 μm, bacterial length <2.5 μm, F_*b*_ (force on the bond) < 0.113 pN, velocity >175 μm/s, and trajectories with fewer than three successive interactions/track (**Fig. S4**).

### Intravenous (IV) perfusion experiments

#### Intravenous infection and perfusion

These experiments were performed largely as described by Caine et al. ^16^ Briefly, ten mice per experimental group (90 total across 9 groups) were inoculated via tail vein with 1 × 10^8^ bacteria washed twice in ice-cold PBS−/−, resuspended to 1 × 10^9^ /ml in PBS and injected using a 29-gauge needle (BD, Mississauga, ON, Canada) after warming to RT. At 1 h, 1 day (24 h) or 1 week, mice were anesthetized with 1% isoflurane or 100 mg ketamine/kg (Rogar/STB) of body weight and 10 mg/kg xylazine (MTC Pharmaceuticals), delivered by intraperitoneal injection. Anesthesia was confirmed by toe pinch, the heart was exposed, and 100 μl of blood was collected using a small cut in the right atrium. A 24Gx.75in catheter (BD) was inserted into the left ventricle and mice were perfused with 20 ml 0.9% sterile sodium chloride (NaCl) at 4 ml/min. Successful perfusion was confirmed by blanching of tissues. Whole bladder, ventral abdominal skin sections and right hemi-sections of heart, brain and patella were rinsed with 1x PBS−/−, blotted dry on tissue, snap-frozen on dry ice, then stored at −80 °C.

#### DNA extraction and qPCR measurement of bacterial and mouse DNA copy number

Total genomic DNA was isolated from tissues using a PureLink Genomic DNA Mini Kit (Fisher) and collected in 100 μl kit elution buffer. Samples for which extraction was unsuccessful were excluded from further analysis. These were identified as samples with DNA concentration values <10% of the median DNA concentration for the same tissue from the same experimental group (<0.1% of samples). qPCR amplification of *flaB* and *nido* sequences was performed in technical sextuplicate and triplicate, respectively, as described ^56,57^, except *nido* PCR conditions were: Step 1: 95 °C 5 min; Step 2: 50 cycles 98 °C 6 s, 60 °C 3 s, 68 °C 5 s; Step 3 melt curve analysis from 60-95 °C. Measurements with abnormal amplification curves, melting temperature or cycle threshold values more than 1.5 standard deviations from the mean for each biological replicate were excluded from subsequent analysis. Median *flaB* and *nido* copy number values for each sample were calculated from remaining values and standard curve values obtained for serial dilutions of known quantities of plasmids bearing *flaB* and *nido* sequences.

#### Clearance-adjusted measurement of tissue-specific dissemination

To control for potential effects of differences in bacterial clearance among experimental groups, differences in tissue cellularity and resulting nido values among tissues, and differences in the lowest detectable limit for *flaB*:*nido* ratios for tissues with distinct host cell densities, we calculated the median *nido* value for the same tissue from all experimental groups from all experiments, and multiplied the reciprocal of this value by the smallest number of *flaB* copies detectable by qPCR (0.17, or 1 copy detected per six technical replicates) to obtain the zero threshold *flaB*:*nido* for each tissue. To normalize the zero threshold for all tissues to the same value and adjust all *flaB*:*nido* values for tissue-specific differences in cell density, we calculated the zero threshold value for each tissue relative to the tissue with the lowest cell density (blood) then multiplied the *flaB*:*nido* values for samples from each tissue by the normalization value for that tissue compared to blood. All *flaB*:*nido* ratios for all tissues less than the universal zero threshold value (the threshold value for blood) were converted to the blood zero threshold value.

Non-zero *flaB*:*nido* ratios that were 5-fold greater or less than 20% of the median value for the same experimental group and tissue were respectively converted to 5-fold maxima and 20% minima calculated using the median value for that group and tissue. We then calculated the sum of the *flaB*:*nido* ratios for all tissues in each mouse (referred to as “All” in **Fig. 2A**), identified mice with suboptimal tail vein injections (<10% of the median of sums of *flaB*:*nido* ratios in all tissues for each mouse in the same experimental group; <1% of all mice in all experiments), and excluded these mice from subsequent analyses.

Finally, the percentage of total bacterial burden for each mouse localized to each tissue was calculated by comparing the tissue *flaB*:*nido* ratio to the sum of *flaB*:*nido* ratios for all tissues from the same mouse. To compare experimental group effect sizes to control groups, the percent total value for the same tissue from each mouse was expressed as a fold difference compared to the median percent total value for the same tissue in control mice.

## Acknowledgements

Critical review of manuscript: Moriarty lab members. Technical support: Peter Gilgan Centre for Research Live Cell Imaging Facility; University of Toronto Division of Comparative Medicine.

## SUPPLEMENTAL TABLES

**Table S1:**
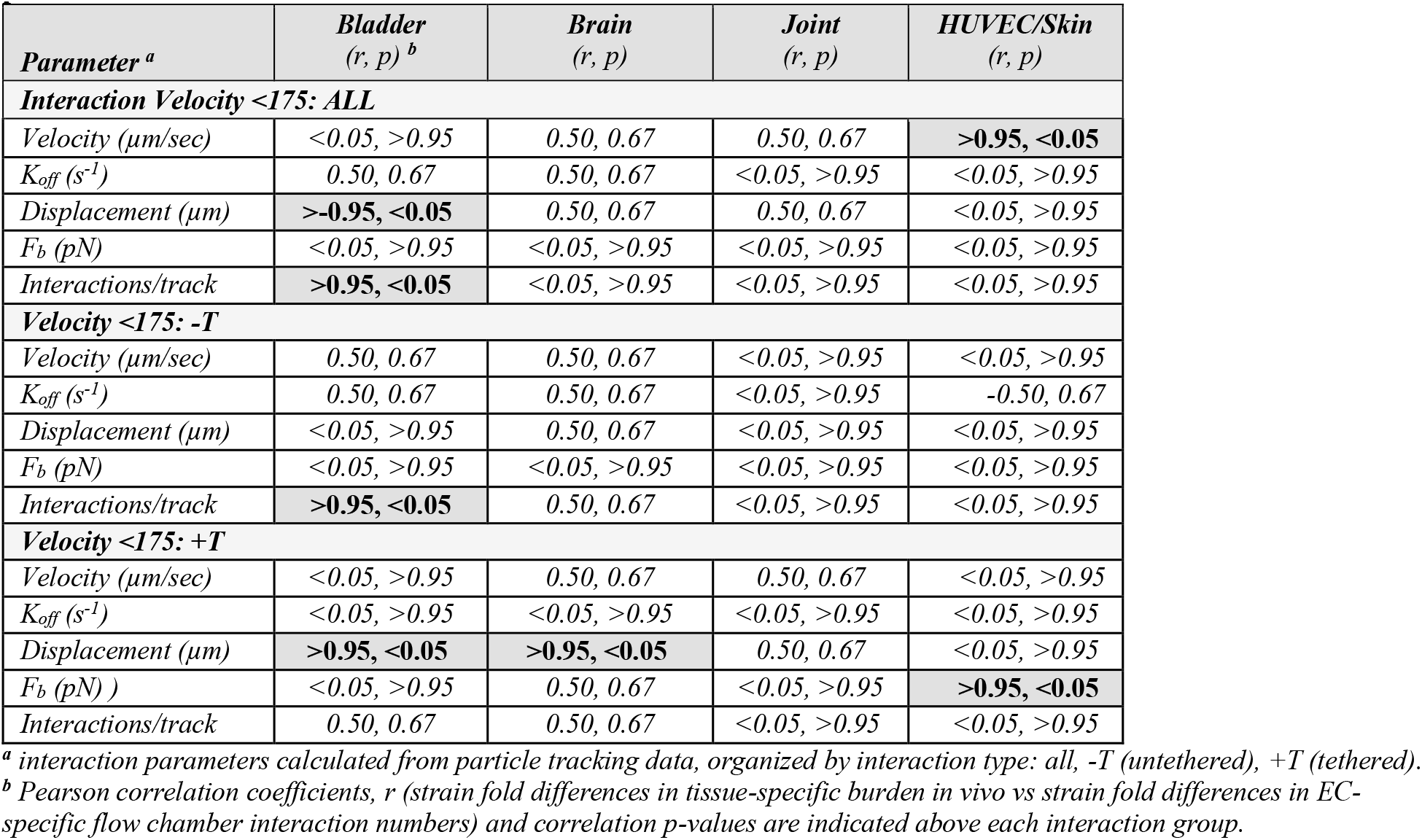
Correlation analysis: tissue burden *in vivo* vs flow chamber interaction parameters, 1 h, 1 d, 1 w post-IV inoculation.

| Parameter <sup>a</sup> | Bladder<br>(*r*, *p*) <sup>b</sup> | Brain<br>(*r*, *p*) | Joint<br>(*r*, *p*) | HUVEC/Skin<br>(*r*, *p*) |
| --- | --- | --- | --- | --- |
| **Interaction Velocity <175: ALL** |  |  |  |  |
| Velocity (μm/sec) | <0.05, >0.95 | 0.50, 0.67 | 0.50, 0.67 | **>0.95, <0.05** |
| *K<sub>off</sub>* (s<sup>-1</sup>) | 0.50, 0.67 | 0.50, 0.67 | <0.05, >0.95 | <0.05, >0.95 |
| Displacement (μm) | **>-0.95, <0.05** | 0.50, 0.67 | 0.50, 0.67 | <0.05, >0.95 |
| *F<sub>b</sub>* (pN) | <0.05, >0.95 | <0.05, >0.95 | <0.05, >0.95 | <0.05, >0.95 |
| Interactions/track | **>0.95, <0.05** | <0.05, >0.95 | <0.05, >0.95 | <0.05, >0.95 |
| **Velocity <175: -T** |  |  |  |  |
| Velocity (μm/sec) | 0.50, 0.67 | 0.50, 0.67 | <0.05, >0.95 | <0.05, >0.95 |
| *K<sub>off</sub>* (s<sup>-1</sup>) | 0.50, 0.67 | 0.50, 0.67 | <0.05, >0.95 | -0.50, 0.67 |
| Displacement (μm) | <0.05, >0.95 | 0.50, 0.67 | <0.05, >0.95 | <0.05, >0.95 |
| *F<sub>b</sub>* (pN) | <0.05, >0.95 | <0.05, >0.95 | <0.05, >0.95 | <0.05, >0.95 |
| Interactions/track | **>0.95, <0.05** | 0.50, 0.67 | <0.05, >0.95 | <0.05, >0.95 |
| **Velocity <175: +T** |  |  |  |  |
| Velocity (μm/sec) | <0.05, >0.95 | 0.50, 0.67 | 0.50, 0.67 | <0.05, >0.95 |
| *K<sub>off</sub>* (s<sup>-1</sup>) | <0.05, >0.95 | <0.05, >0.95 | <0.05, >0.95 | <0.05, >0.95 |
| Displacement (μm) | **>0.95, <0.05** | **>0.95, <0.05** | 0.50, 0.67 | <0.05, >0.95 |
| *F<sub>b</sub>* (pN) | <0.05, >0.95 | 0.50, 0.67 | <0.05, >0.95 | **>0.95, <0.05** |
| Interactions/track | 0.50, 0.67 | 0.50, 0.67 | <0.05, >0.95 | <0.05, >0.95 |

<sup>a</sup> interaction parameters calculated from particle tracking data, organized by interaction type: all, -T (untethered), +T (tethered).
<sup>b</sup> Pearson correlation coefficients, *r* (strain fold differences in tissue-specific burden *in vivo* vs strain fold differences in EC-specific flow chamber interaction numbers) and correlation *p*-values are indicated above each interaction group.

**Table S2:** Primary human endothelial cell types used in this study.

| Vascular bed | Cell Type | Abbreviation | Single or multiple donors <sup>a</sup> | Sex | Source | Catalog number |
| --- | --- | --- | --- | --- | --- | --- |
| Aorta | arterial | HAEC | single | female | Lonza Inc | CC-2535 |
| Bladder | microvascular | HMVEC- Bd | single | female | Lonza Inc | CC-7016 |
| Brain cortex | microvascular | HMVEC-Br | single | male | Cell Systems | ACBRI 376 |
| Synovial tissue | microvascular | HMVEC-Joint | single | female | Cell Systems | ACBRI 488 |
| Umbilical cord vein | venous | HUVEC | multiple | female | Lonza Inc | CC-2519 |

<sup>a</sup> all endothelia were derived from healthy donors.

**Table S3:**
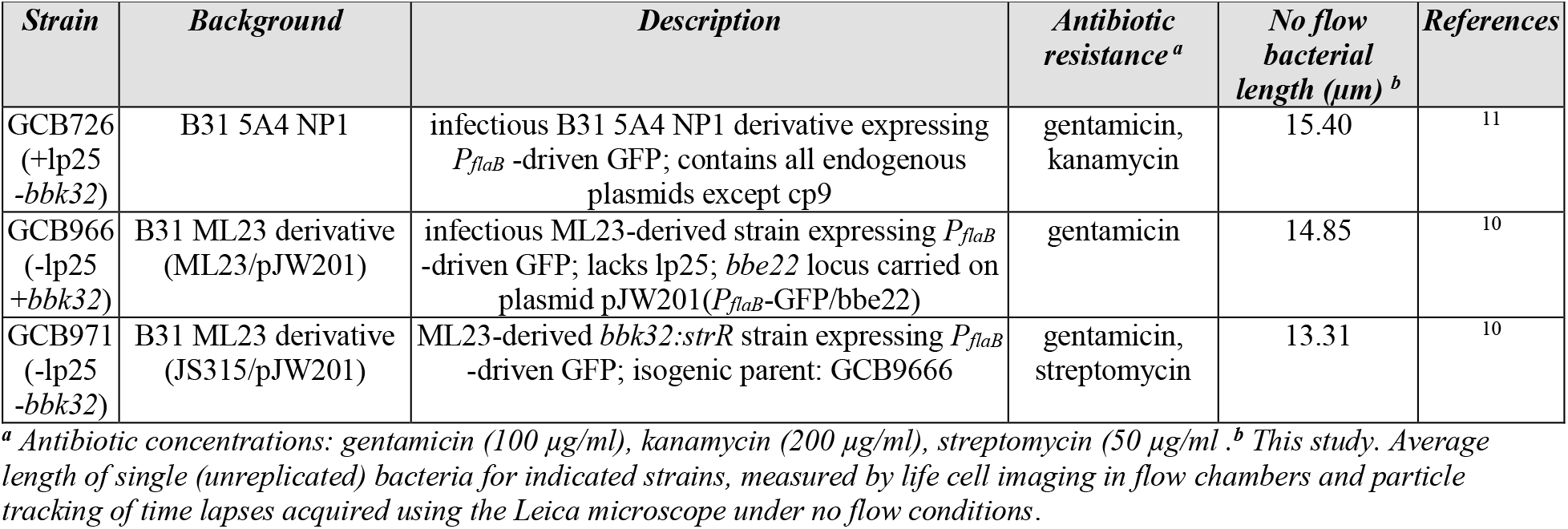
*B. burgdorferi* strains used in this study.

**Table S4:** Particle tracking conditions.

| *Protocol* | *Parameters* | **GCB726** | **GCB966** | **GCB971** |
| --- | --- | --- | --- | --- |
| A | **Object size** ($\mu\text{m}^2$): 1.5-1.1X no-flow bacterial length (BL) | 1.5-16.95 | 1.5-16.33 | 1.5-14.64 |
| | **Object Skeletal length** ($\mu\text{m}$): 2.5-1X BL | 2.5- 15.4 | 2.5- 14.85 | 2.5- 13.31 |
|  | **Object Intensity** | 17-255 |  |  |
|  | **Object joining (tracking) algorithm** | Trajectory Variation (0.01), Estimate Max Distance between Objects Automatically |  |  |
| | **Maximum Track Velocity** ($\mu\text{m/s}$) | 300 | | |
| B | **Object size** ($\mu\text{m}^2$): 1.5-1.1X no-flow bacterial length (BL) | 1.2-53.28 | 1.2-51.38 | 1.2-46.05 |
| | **Object Skeletal length** ($\mu\text{m}$): 2.5-1X BL | 1.4-15.4 | 1.4-14.85 | 1.4-13.31 |
|  | **Object Intensity** | 40-135 |  |  |
|  | **Object joining (tracking) algorithm** | See details above in protocol A |  |  |
| | **Maximum Track Velocity** ($\mu\text{m/s}$) | $\leq 185 \mu\text{m/sec}$ | | |
| C | **Object size** ($\mu\text{m}^2$): 1.5-1.1X no-flow bacterial length (BL) | 5.0-53.28 | 5.0-51.38 | 5.0-46.05 |
| | **Object Skeletal length** ($\mu\text{m}$): 2.5-1X BL | 1.4-15.4 | 1.4-14.85 | 1.4-13.31 |
|  | **Object Intensity** | 40-135 |  |  |
|  | **Object joining (tracking) algorithm** | See details above in protocol A |  |  |
| | **Maximum Track Velocity** ($\mu\text{m/s}$) | $\leq 300 \mu\text{m/sec}$ | | |
| D | **Object size** ($\mu\text{m}^2$): 1.5-1.1X no-flow bacterial length (BL) | 1.2-53.28 | 1.2-51.38 | 1.2-46.05 |
| | **Object Skeletal length** ($\mu\text{m}$): 2.5-1X BL | 1.4-15.4 | 1.4-14.85 | 1.4-13.31 |
|  | **Object Intensity** | 110-255 |  |  |
|  | **Object joining (tracking) algorithm** | See details above in protocol A |  |  |
| | **Maximum Track Velocity** ($\mu\text{m/s}$) | $\leq 185 \mu\text{m/sec}$ | | |
| E | **Object size** ($\mu\text{m}^2$): 1.5-1.1X no-flow bacterial length (BL) | 5.0-53.28 | 5.0-51.38 | 5.0-46.05 |
| | **Object Skeletal length** ($\mu\text{m}$): 2.5-1X BL | 1.4-15.4 | 1.4-14.85 | 1.4-13.31 |
|  | **Object Intensity** | 110-255 |  |  |
|  | **Object joining (tracking) algorithm** | See details above in protocol A |  |  |
| | **Maximum Track Velocity** ($\mu\text{m/s}$) | $\leq 300 \mu\text{m/sec}$ | | |
| F | **Object size** ($\mu\text{m}^2$): 1.5-1.1X no-flow bacterial length (BL) | 1.5-16.95 | 1.5-16.33 | 1.5-14.64 |
| | **Object Skeletal length** ($\mu\text{m}$): 1.4-1X BL | 1.4-15.4 | 1.4-14.85 | 1.4-13.31 |
|  | **Object Intensity** | 17-255 |  |  |
| | **Object joining (tracking) algorithm** | Shortest Path (Max Distance 15 $\mu\text{m}$) | | |
| | **Maximum Track Velocity** ($\mu\text{m/s}$) | 200 | | |

## SUPPLEMENTAL FIGURES

**Fig S1:**
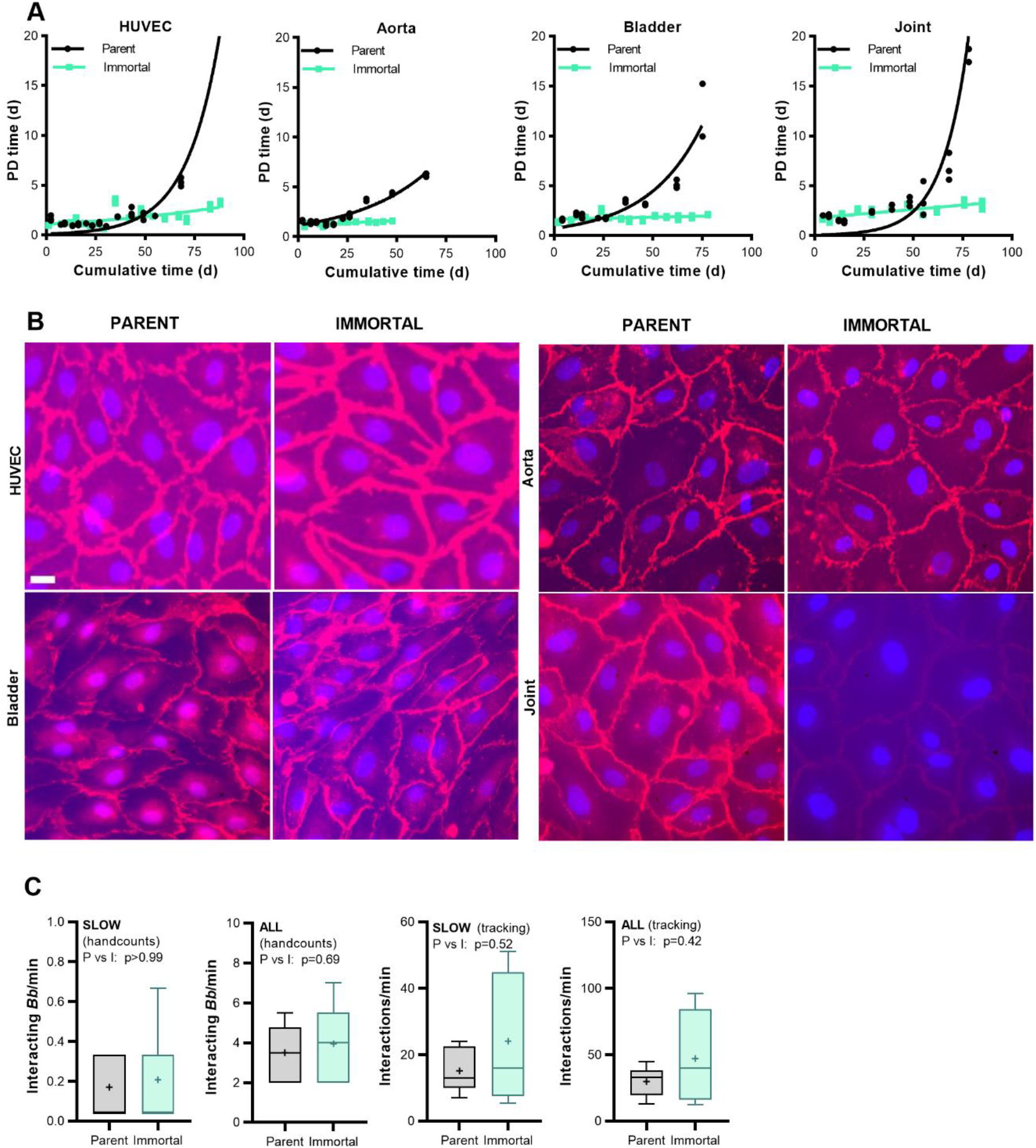
Population doubling times and VE-cadherin expression in parent and immortalized ECs. **(A)** Population doubling (PD) time plots and non-linear regressions for parent (P) and hTERT-immortalized (I) ECs. **(B)** IF visualization of VE-cadherin (red) in P and I ECs counterstained with DAPI (blue). Scale bar: 30 μm. **(C)** Data are shown as Tukey box and whiskers plots of total (all) and slow (velocity <125 μm/s) interaction numbers under flow for all bacterial strains (+lp25+*bbk32,* −lp25+*bbk32,* −lp25-*bbk32*) and ECs derived from aorta, bladder and joint, measured by manual and particle tracking enumeration (hand counts, tracking, respectively). For all Fig. S1 experiments, N ≥ 3 independent EC and bacterial cultures. Statistics: Mann-Whitney or Welch’s t-test (depending on normality of distribution, determined by normality tests) of log-transformed data.

**Fig. S2:**
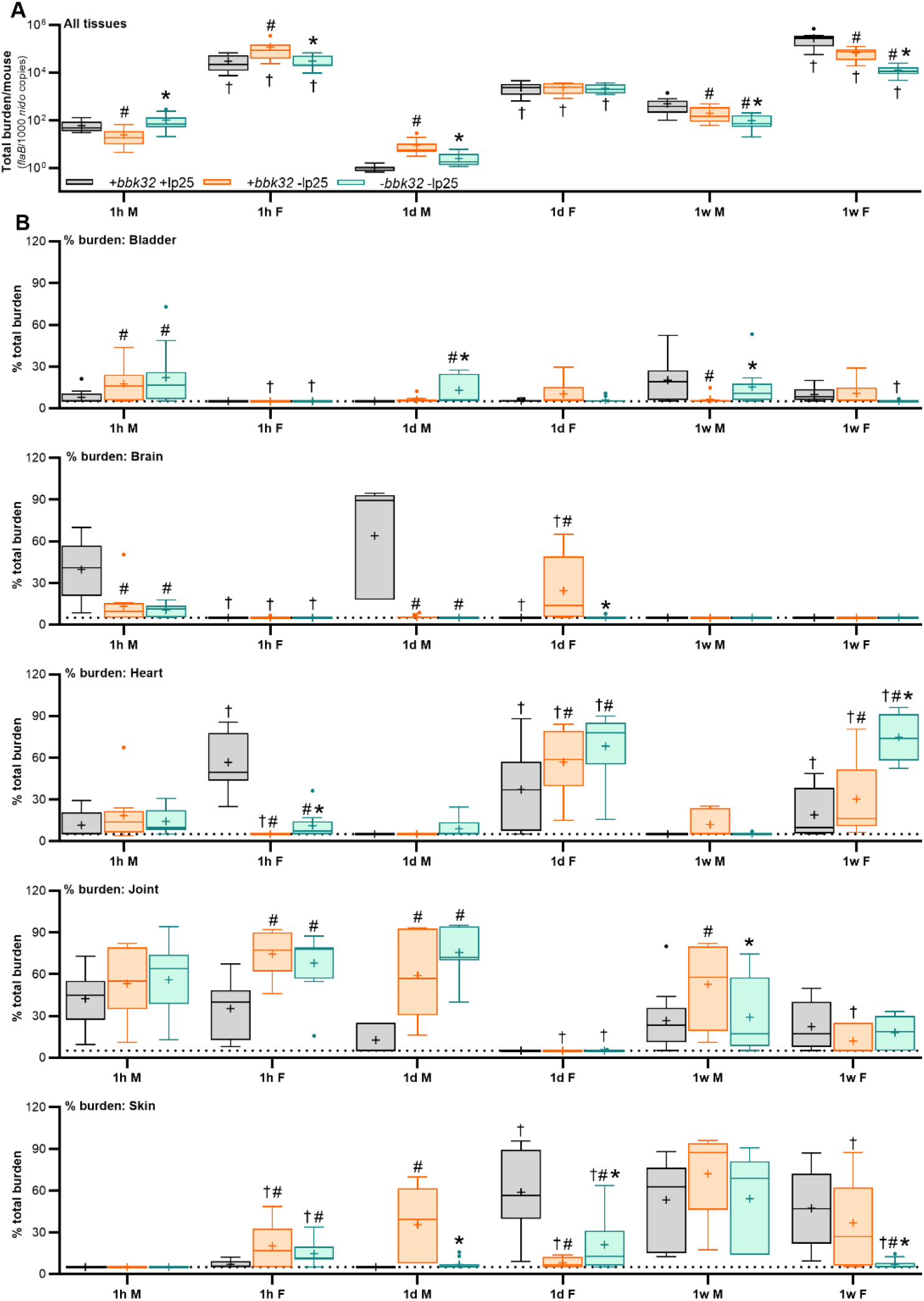
*B. burgdorferi* dissemination to tissues after intravenous inoculation of male or female mice. Total *B. burgdorferi* burden in all tissues **(A)** and percent of total burden in indicated tissues at 1 h, 1day (d) and 1 week (w) after intravenous inoculation of female and male mice **(B)**, measured by quantitative PCR. Data are shown as Tukey box and whiskers plots. N≥26 mice/experimental group. Statistics: two-way ANOVA of log-transformed values, with Holm-Sidak post-tests. * indicates p<0.05 vs −lp25-*bbk32*. # indicates p<0.05 vs +lp25+*bbk32*.

**Fig. S3:**
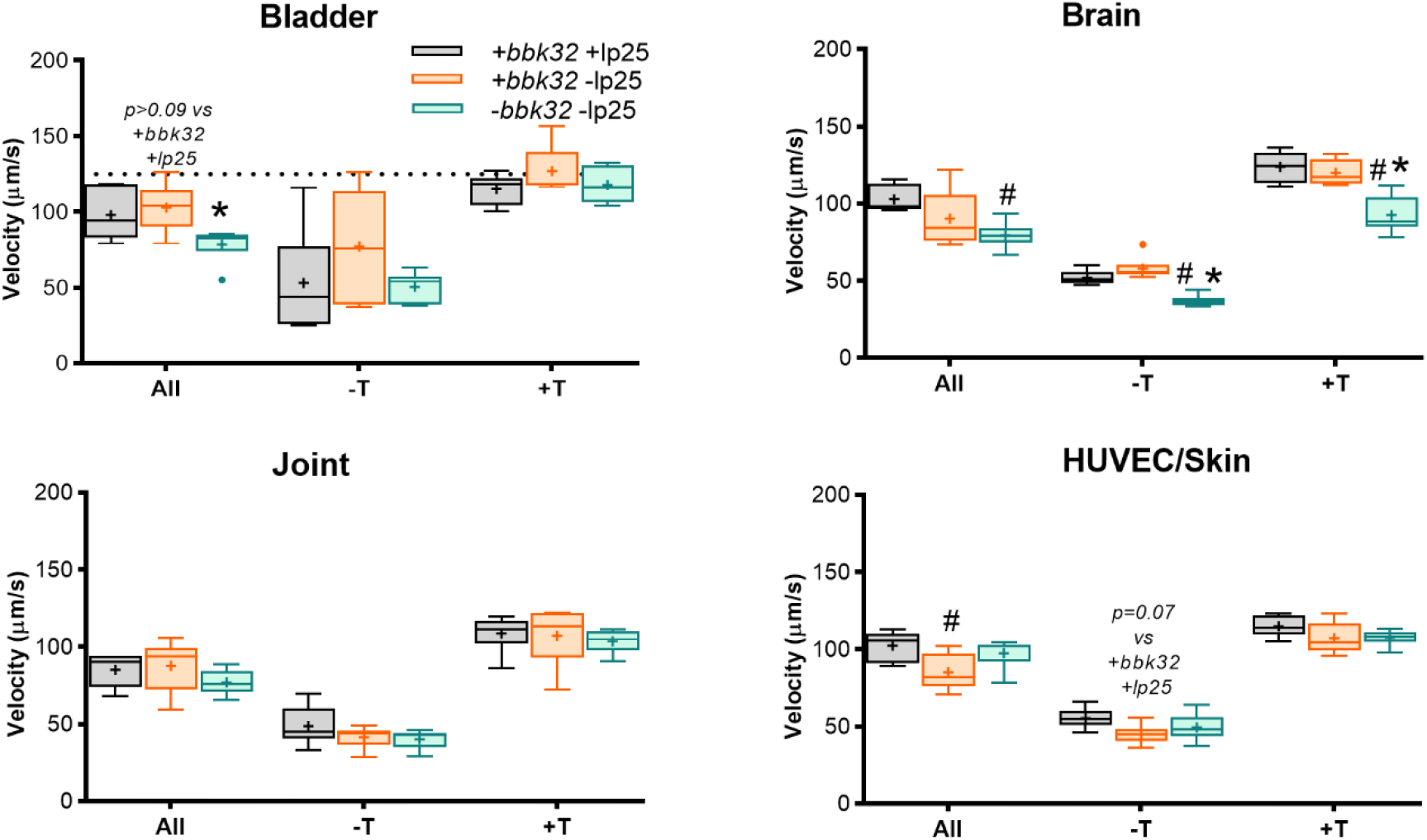
Bacterial-endothelial interaction velocity in flow chambers predict dissemination patterns *in vivo*. *B. burgdorferi*-EC interaction velocity (μm/s:) for all, −T (untethered) and +T (tethered) interactions in flow chambers, measured by particle tracking. Data shown as Tukey box and whiskers plots. N ≥3 biological replicates/experimental group. Statistical comparisons within each interaction group: ANOVA with Tukey’s post-tests. * indicates p<0.05 vs −lp25+*bbk32*. # indicates p<0.05 vs +lp25+*bbk32*. Pearson correlation coefficients, *r* (strain fold differences in tissue-specific burden *in vivo* vs strain fold differences in EC-specific flow chamber interaction numbers) and correlation *p*-values are indicated above each interaction group. Correlation values for all other interaction parameters are provided in **Tables 1 and S1**.

**Fig. S4:**
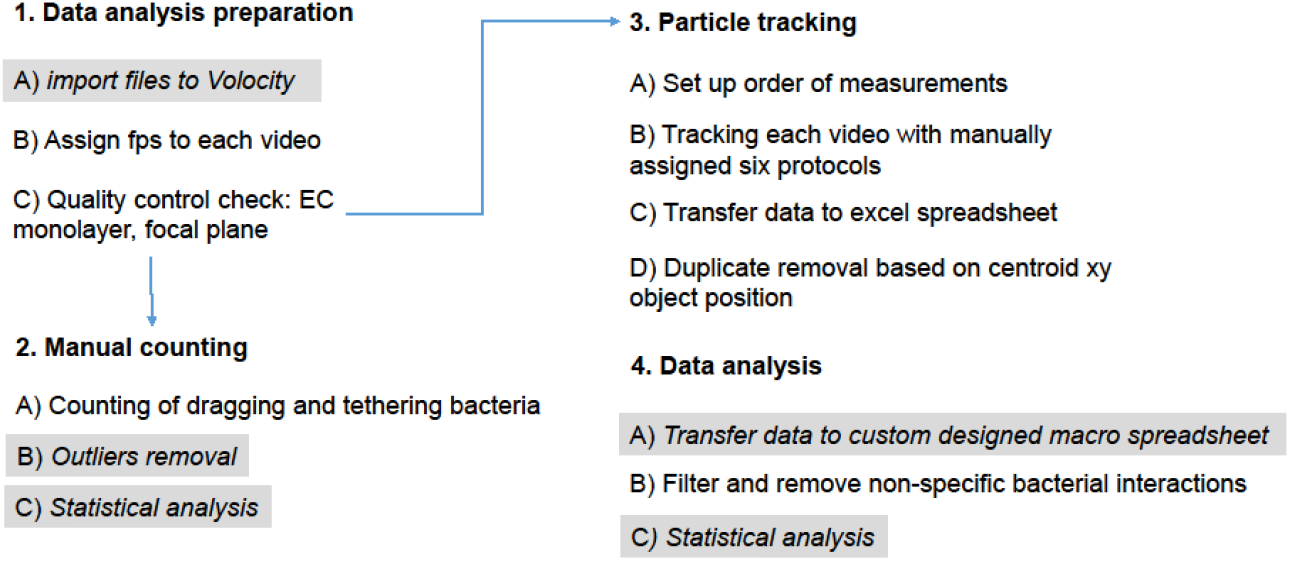
Bioinformatics summary of particle tracking and measuring biophysical properties of *B. burgdorferi*-endothelial interactions under flow. Quality control and analysis steps in the time lapse bioinformatics. Highlighted in grey are steps performed using automated spreadsheets and software (Volocity, GraphPad Prism). Other steps are performed manually.

